# Reprogramming of the FOXA1 cistrome in treatment-emergent neuroendocrine prostate cancer

**DOI:** 10.1101/2020.10.23.350793

**Authors:** Sylvan C. Baca, David Y. Takeda, Ji-Heui Seo, Justin Hwang, Sheng Yu Ku, Rand Arafeh, Taylor Arnoff, Supreet Agarwal, Connor Bell, Edward O’Connor, Xintao Qiu, Sarah Abou Alaiwi, Rosario I. Corona, Marcos A. S. Fonseca, Claudia Giambartolomei, Paloma Cejas, Klothilda Lim, Monica He, Anjali Sheahan, Amin Nassar, Jacob E. Berchuck, Lisha Brown, Holly M. Nguyen, Ilsa M. Coleman, Arja Kaipainen, Navonil De Sarkar, Peter S. Nelson, Colm Morrissey, Keegan Korthauer, Mark M. Pomerantz, Leigh Ellis, Bogdan Pasaniuc, Kate Lawrenson, Kathleen Kelly, Amina Zoubeidi, William C. Hahn, Himisha Beltran, Henry W. Long, Myles Brown, Eva Corey, Matthew L. Freedman

## Abstract

Lineage plasticity, the ability of a cell to alter its identity, is an increasingly common mechanism of adaptive resistance to targeted therapy in cancer^1,2^. An archetypal example is the development of neuroendocrine prostate cancer (NEPC) after treatment of prostate adenocarcinoma (PRAD) with inhibitors of androgen signaling. NEPC is an aggressive variant of prostate cancer that aberrantly expresses genes characteristic of neuroendocrine (NE) tissues and no longer depends on androgens. To investigate the epigenomic basis of this resistance mechanism, we profiled histone modifications in NEPC and PRAD patient-derived xenografts (PDXs) using chromatin immunoprecipitation and sequencing (ChIP-seq). We identified a vast network of *cis*-regulatory elements (N~15,000) that are recurrently activated in NEPC. The FOXA1 transcription factor (TF), which pioneers androgen receptor (AR) chromatin binding in the prostate epithelium^3,4^, is reprogrammed to NE-specific regulatory elements in NEPC. Despite loss of dependence upon AR, NEPC maintains FOXA1 expression and requires FOXA1 for proliferation and expression of NE lineage-defining genes. Ectopic expression of the NE lineage TFs ASCL1 and NKX2-1 in PRAD cells reprograms FOXA1 to bind to NE regulatory elements and induces enhancer activity as evidenced by histone modifications at these sites. Our data establish the importance of FOXA1 in NEPC and provide a principled approach to identifying novel cancer dependencies through epigenomic profiling.

## Introduction

In recent years, potent AR pathway inhibitors have extended the survival of patients with metastatic prostate cancer^5,6^. Prostate tumors inevitably escape AR inhibition through reactivation of AR signaling or, increasingly, via lineage plasticity^1,7^. The mechanisms underlying lineage plasticity remain unclear but likely involve transdifferentiation of PRAD to NEPC rather than *de novo* emergence of NEPC. NEPC and PRAD tumors from an individual patient share many somatic DNA alterations, implying a common ancestral tumor clone^8^. While the genomic profiles of NEPC and PRAD are relatively similar, their gene expression profiles and clinical behavior differ markedly^9^. We therefore set out to characterize epigenomic differences between NEPC and PRAD, hypothesizing that reprogramming of distinct regulatory elements drives their divergent phenotypes.

## Results

We performed ChIP-seq for the histone post-translational modification H3K27ac to identify active regulatory elements in the LuCaP PDX series^10^, a set of xenografts derived from advanced PRAD (N=22) and treatment-emergent NEPC (N=5). We identified a median of 55,095 H3K27ac peaks per sample (range 37,599-74,640) (Supplementary Table 1). Notably, the transcriptomes of the LuCaP PDXs reflect differences in gene expression observed between clinical PRAD and NEPC metastases (Supplementary Fig. 1a), indicating their relevance to clinical prostate cancer.

Unsupervised hierarchical clustering and principal component analysis based on genome-wide H3K27 acetylation cleanly partitioned NEPC and PRAD LuCaP PDXs (Fig. 1a, Supplementary Fig. 1b, c). We identified 14,985 sites with eight-fold or greater increases in H3K27 acetylation in NEPC compared to PRAD at an adjusted *p*-value of 10^-3^. We termed these sites neuroendocrine-enriched candidate regulatory elements (“Ne-CREs”; Fig. 1b, Supplementary Table 2, Supplementary Fig. 1d). A smaller set of sites (4,338) bore greater H3K27ac signal in PRAD (termed “Ad-CREs”). Liver metastases from clinical NEPC and PRAD demonstrated enrichment of H3K27ac at Ne-CREs and Ad-CREs, respectively, confirming that the LuCaP PDX models reflect lineage-specific epigenomic features of clinical prostate tumors (Supplementary Fig. 1e).

**Figure 1.**
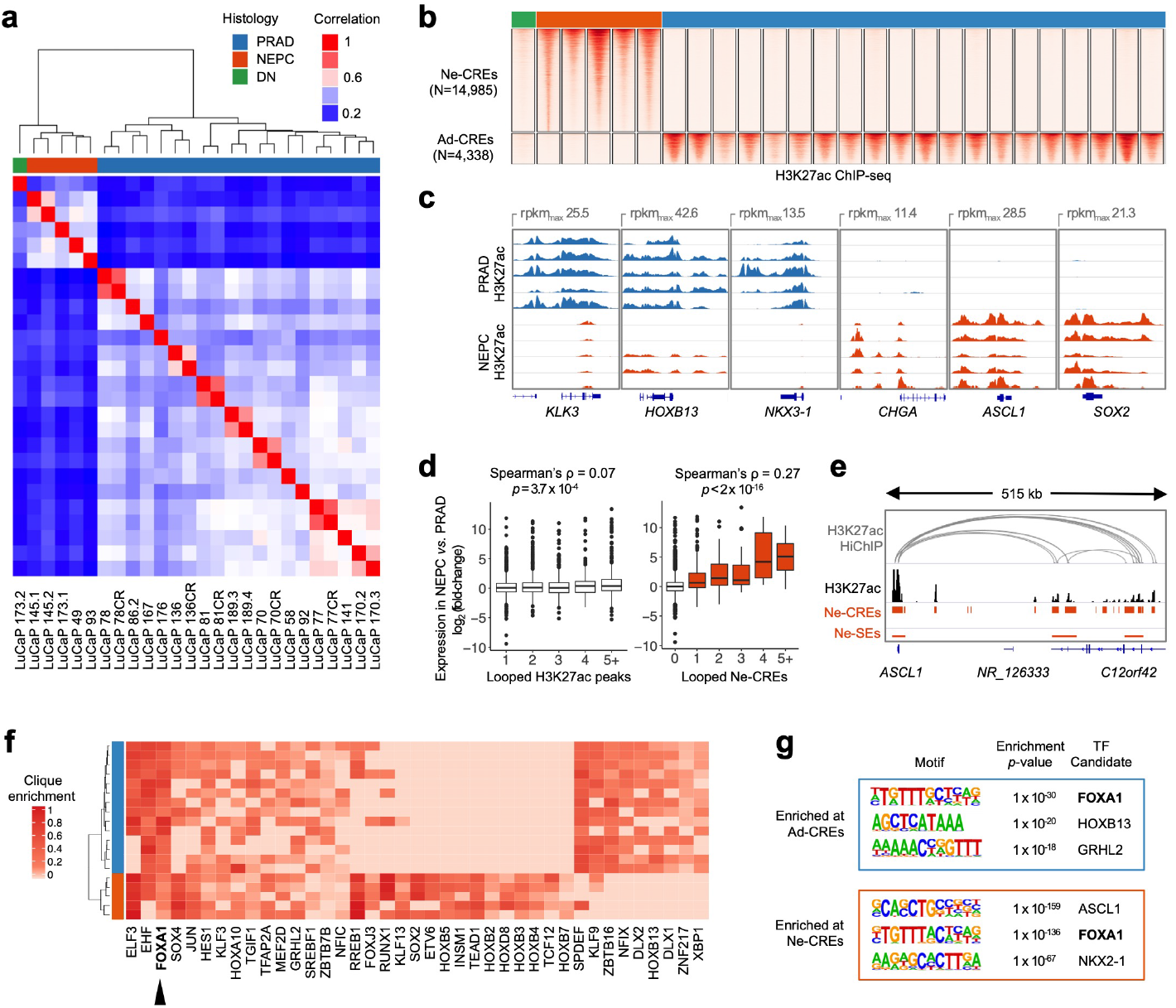
Epigenomic divergence of PRAD and NEPC. **a**, Hierarchical clustering of PRAD and NEPC based on sample-to-sample correlation of H3K27ac profiles. “DN” (“double-negative”) indicates a LuCaP PDX without AR or NE marker expression (see also Supplementary Figure 1). **b**, Heatmaps of normalized H3K27ac tag densities at differentially H3K27-acetylated regions (±2kb from peak center) between NEPC and PRAD. “CREs” signify candidate regulatory elements. **c**, H3K27ac signal near selected prostate-lineage and NEPC genes. Five representative samples from each histology are shown. **d**, Differential expression (NEPC *vs*. PRAD) of genes with the indicated number of distinct looped H3K27ac peaks (left) or Ne-CREs (right) detected by H3K27ac HiChIP in LuCaP 173.1 (NEPC). Wilcoxon *p*-value is indicated for comparison of genes with loops to one Ne-CRE or H3K27ac peak versus two or more. **e**, H3K27ac HiChIP loops in LuCaP 173.1 from *ASCL1* to Ne-CREs and NEPC-restricted super-enhancers (Ne-SEs). H3K27ac tag density for LuCaP 173.1 is shown in black. **f**, Candidate master transcription factors in NEPC and PRAD based on regulatory clique enrichment (see methods). **g**, Three most significantly enriched nucleotide motifs present in >10% of Ad-CREs or Ne-CREs by *de novo* motif analysis.

Ad-CREs were found near prostate lineage genes such as *KLK3, HOXB13,* and *NKX3-1*,x while Ne-CREs resided near genes enriched for neuronal and developmental annotations, including *CHGA, ASCL1,* and *SOX2^11^* (Fig. 1c, Supplementary Table 3). Genes with higher expression in NEPC compared to PRAD were enriched for nearby Ne-CREs (Supplementary Fig. 1f) and formed three-dimensional contacts with a greater number of Ne-CREs as assessed by H3K27ac HiChIP (Fig. 1d, Supplementary Fig. 1g-h, and Supplementary Tables 4 and 5). For example, *ASCL1,* which encodes a neural lineage TF that is highly upregulated in NEPC (Supplementary Fig. 1a), interacts with 15 gene-distal Ne-CREs between 280kb and 465kb telomeric to *ASCL1,* including two novel NEPC-restricted super-enhancers within intronic regions of *C12ORF42* (Fig. 1e). These results suggest that Ne-CREs regulate neuroendocrine transcriptional programs through interaction with NEPC gene promoters.

We nominated candidate TFs that may orchestrate NEPC lineage gene expression by binding to Ne-CREs. Lineage-defining TF genes often reside within densely H3K27-acetylated super-enhancers^12^ and form core regulatory circuits, or “cliques”, by mutual binding of one another’s *cis*-regulatory regions^13,14^. Several TFs showed clique enrichment specifically in NEPC (Fig. 1f) and/or were encompassed by NEPC-restricted super-enhancers (Supplementary Fig. 2), including known NE lineage TFs *(e.g., ASCL1* and *INSM1)* and novel candidates such as *HOXB2-5.*

Notably, a single TF gene, *FOXA1,* demonstrated clique enrichment in all NEPC and PRAD LuCaP PDXs (Fig. 1f). FOXA1 is a pioneer TF of endodermal tissues^3^ with a critical role in prostate development^4^ but no characterized function in NEPC. The forkhead motif recognized by FOXA1 was the second most significantly enriched nucleotide sequence within Ne-CREs (Fig. 1g). FOXA2, a previously-reported NEPC TF^15^, does not wholly account for the forkhead motif enrichment because FOXA2 was not expressed in several NEPC samples (Figs. 2a,b; Supplementary Fig. 3a). In contrast, FOXA1 was expressed in all NEPCs (Fig 2a-b; Table S6) as well as in resident neuroendocrine cells of benign prostate tissue (Supplementary Fig. 3).

**Figure 2.**
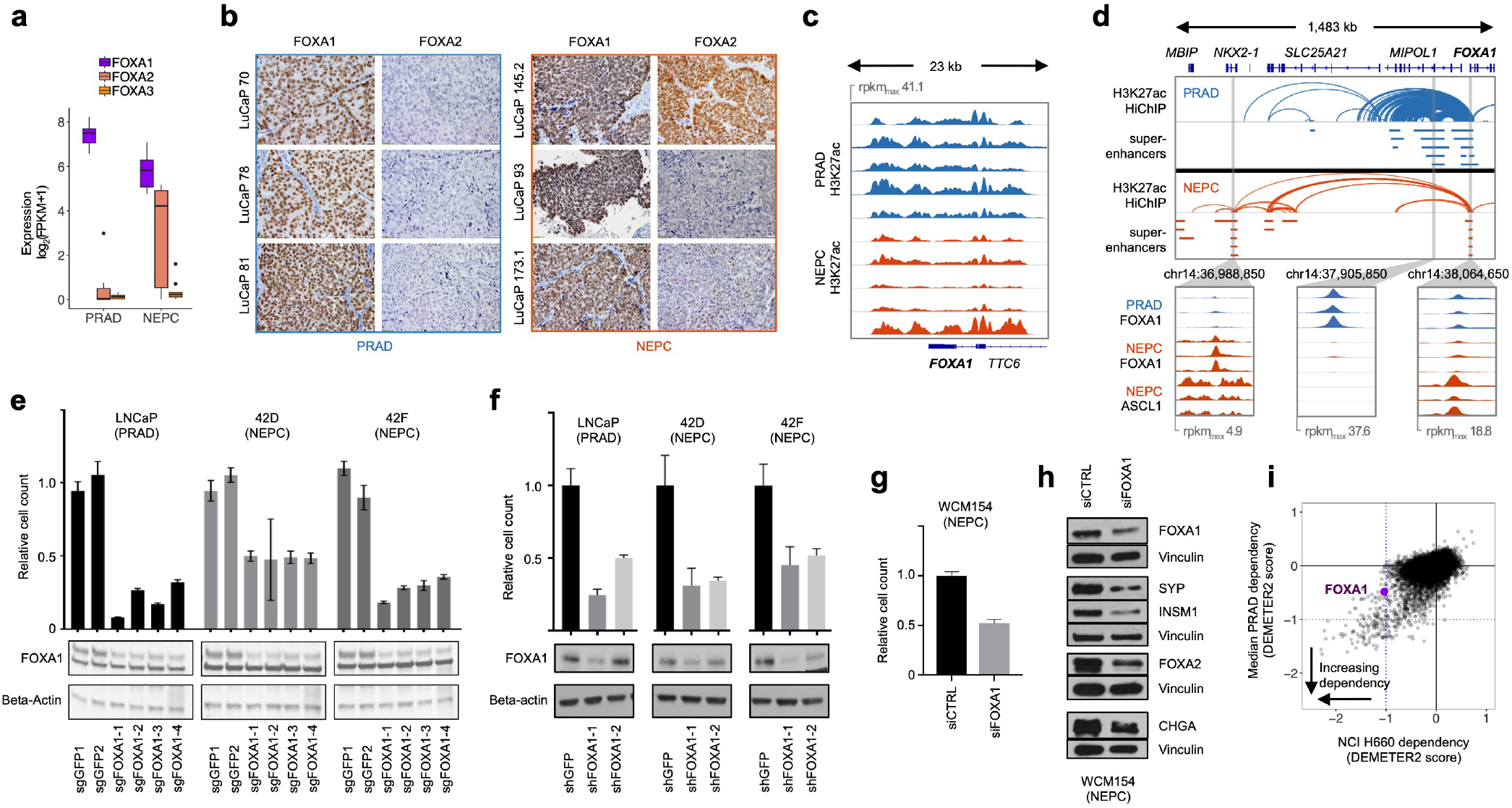
FOXA1 remains a critical lineage transcription factor in NEPC. **a**, Transcript expression of FOXA family TFs in LuCaPs PDXs (5 NEPC and 5 PRAD; 2 replicates each). **b**, FOXA1/FOXA2 immunohistochemistry in six representative PDXs. **c**, H3K27ac profiles at *FOXA1* in five representative PRAD and NEPC PDXs. **d**, H3K27ac HiChIP loops near *FOXA1* in LuCaP 173.1 (NEPC) and LNCaP (PRAD). Bars indicate super-enhancers in five representative LuCaPs of each lineage. Blowups show ChIP-seq read pileups for FOXA1 and ASCL1 in PDXs of the indicated lineage. **e-f**, Proliferation of LNCaP and 42D/42F derivatives with inactivation of FOXA1 by CRISPR (e) or shRNA (f). **g-h**, Proliferation (g) and expression of neuroendocrine marker proteins (h) with siRNA knock-down of FOXA1 in the NEPC organoid model WCM154. **i**, Essentiality of genes in NCI-H660 (NEPC) versus PRAD cell lines in a published shRNA screening dataset^69^. More negative DEMETER2 scores indicate greater dependency. The blue lines indicate the median DEMETER2 score for pan-essential genes.

Multiple lines of investigation supported a pivotal role of FOXA1 in NEPC. A superenhancer encompassed *FOXA1* in all NEPC LuCaP PDXs (Fig. 2c, Supplementary Fig. 2). In NEPC, the *FOXA1* promoter shed contacts with its regulatory region identified in PRAD^16^ and looped to a distinct NEPC-restricted super-enhancer (Fig. 2d). Both the distal superenhancer and promoter were co-bound by FOXA1 and ASCL1, suggesting an auto-regulatory circuit that is characteristic of master transcriptional regulators^17^. Suppression of FOXA1 in a variety of NEPC cellular models^18,19^ demonstrated that FOXA1 is essential for cellular proliferation (Fig. 2e-g) and expression of NE markers, including NE lineage TFs such as FOXA2 and INSM1 (Fig. 2h). Analysis of a published shRNA screen confirmed a dependency on FOXA1 in the NEPC cell line NCI-H660 (Fig. 2i). Thus, FOXA1 exhibits several features of a master transcriptional regulator in NEPC.

We profiled FOXA1 binding sites in NEPC and PRAD using ChIP-seq. FOXA1 relocates to a distinct set of binding sites in NEPC PDXs (Fig. 3a), which overlap with the majority of Ne-CREs (Fig. 3b). In PRAD, Ne-CREs were devoid of FOXA1 binding and heterochromatic as assayed by ATAC-seq, but they acquired FOXA1 binding and chromatin accessibility in NEPC (Fig. 3c). Conversely, Ad-CREs lost FOXA1 binding in NEPC and became less accessible by ATAC-seq. To contextualize the extent of FOXA1 reprogramming in NEPC, we compared FOXA1 binding profiles in normal prostate epithelium, localized PRAD, and PDXs derived from metastatic PRAD. At the same level of stringency, fewer than 500 sites exhibited differential FOXA1 binding between these categories; by comparison, FOXA1 binding was gained at 20,935 and lost at 29,308 sites in NEPC compared to metastatic PRAD (Fig. 3d).

**Figure 3.**
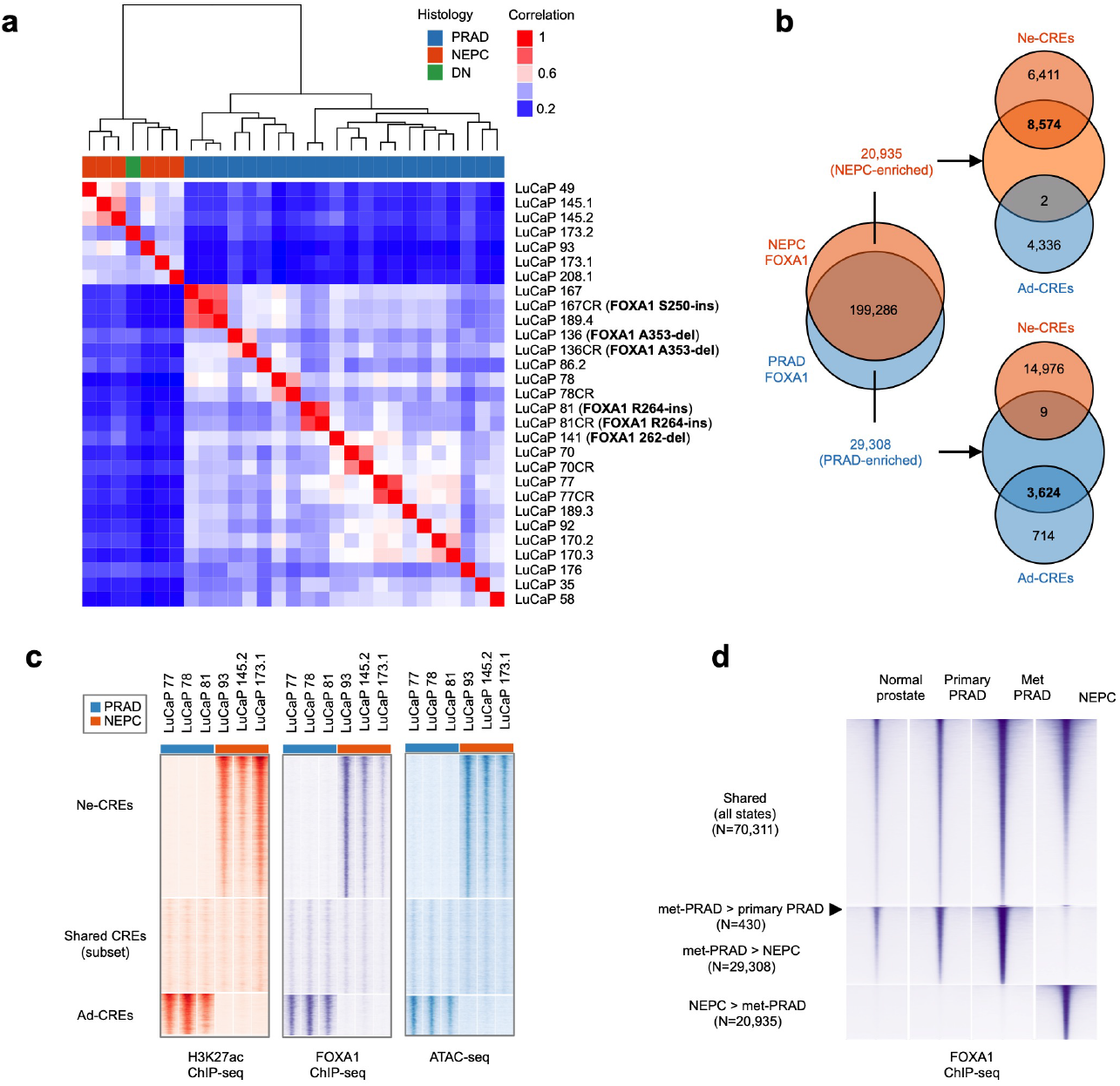
Reprogramming of the FOXA1 cistrome in NEPC. **a**, Hierarchical clustering of LuCaP PDXs by FOXA1 binding profiles. “DN” (“double-negative”) indicates a PDX without AR or NE marker expression. FOXA1 mutational status is noted; see also Table S7) **b**, Venn diagram of lineage-enriched and shared FOXA1 binding sites and their overlap with lineage-enriched candidate regulatory elements (Ad-CREs and Ne-CREs). **c**, Normalized tag densities for H3K27ac/FOXA1 ChIP-seq and ATAC-seq at Ne-CREs and Ad-CREs. Three representative NEPC and PRAD PDXs are shown. **d**, Average normalized tag densities for FOXA1 in normal prostate, primary PRAD, and PDXs derived from PRAD metastases (Met PRAD) or NEPC (five samples in each category) at differential FOXA1 binding sites between these groups. There are insufficient differential sites to display (<100) for the Primary PRAD > Met PRAD comparison and the Primary PRAD *vs*. Normal prostate comparisons.

We sought to understand the mechanism by which FOXA1 binding is reprogrammed in NEPC. In addition to DNA sequence, cooperative binding with partner TFs is an important determinant of pioneer factor localization^20^. Since the motifs recognized by ASCL1 and NKX2-1 were highly enriched at Ne-CREs (Fig. 1g), we tested whether overexpression of these TFs in the PRAD cell line LNCaP could induce FOXA1 binding at Ne-CREs. Overexpression of ASCL1 and NKX2-1 (A+N) increased FOXA1 binding at NEPC-enriched FOXA1 binding sites (Fig. 4a,b) and induced H3K27 acetylation of Ne-CREs (Fig. 4c-f). ASCL1 co-localized with FOXA1 at NEPC-enriched FOXA1 binding sites and Ne-CREs (Fig. 4g-h). A+N expression recapitulated global transcriptional changes between NEPC and PRAD, including suppression of *AR* and induction of *SYP* and *CHGA* (Fig. 4i-k). Thus, ectopic expression of ASCL1 and NKX2-1 is sufficient to partially reprogram FOXA1 binding in PRAD to Ne-CREs and induce *de novo* H3K27 acetylation at these regions, with resultant NEPC gene expression.

**Figure 4.**
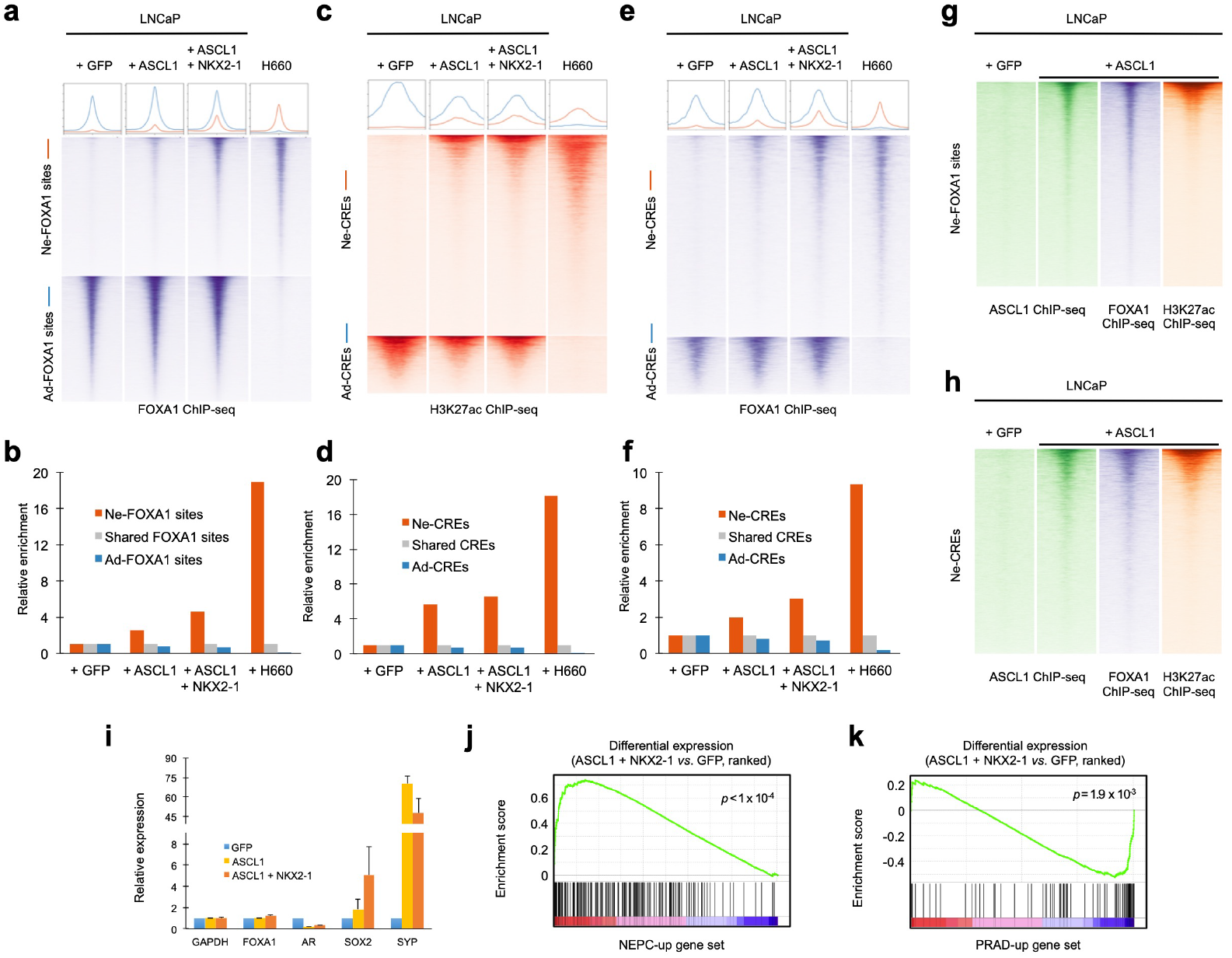
FOXA1 is extensively redistributed at lineage-specific regulatory elements. **a**, Normalized ChIP-seq tag density for FOXA1 at NEPC-enriched and PRAD-enriched FOXA1 binding sites under the indicated conditions. Profile plots (top) represent mean tag density at sites depicted in the heatmaps. **b**, Enrichment of FOXA1 peaks for overlap with NEPC-enriched and PRAD-enriched FOXA1 binding sites in the indicated conditions, normalized to FOXA1 peaks shared between PRAD and NEPC. **c-f**, Normalized ChIP-seq tag density for H3K27ac (c) and FOXA1 (e) at Ne-CREs and Ad-CREs under the indicated experimental conditions. Enrichment of overlap of H3K27ac peaks (d) and FOXA1 peaks (f) with Ne-CREs and Ad-CREs under the indicated conditions. **g-h** Normalized ChIP-seq tag density for ASCL1, FOXA1, and H3K27ac under the indicate experimental conditions at NEPC-enriched FOXA1 sites (g) and Ne-CREs (h). **i**, Effect of ASCL1 overexpression on transcript levels of indicated genes, measured by qPCR. Fold-change relative to +GFP condition is shown, using normalization to GAPDH. The average of three biological replicates is shown for each condition. Error bars represent standard deviation. **j-k**, Gene set enrichment analysis of genes upregulated at least 8-fold in LuCaP NEPC (j) or PRAD (k) at adjusted *p*-value < 10^-18^ Genes are ranked by differential expression between LNCaP + ASCL1 + NKX2-1 and + GFP conditions based on RNA-seq.

Despite intense interest, it remains unclear why PRAD can adopt a seemingly unrelated lineage to overcome androgen blockade, while most cancers do not dramatically alter their cellular identity throughout treatment. Lineage tracing studies have demonstrated that the epithelial cells that give rise to PRAD share a common developmental progenitor with resident neuroendocrine cells in the prostate^21,22^. In this common progenitor cell, Ne-CREs and their FOXA1 binding sites might be physiologically poised for activation upon commitment to a neuroendocrine lineage. In support of this model, genes that are highly expressed in normal neuroendocrine prostate cells are also highly expressed in NEPC (Fig. 5a), and are enriched for nearby Ne-CREs and NEPC-restricted FOXA1 binding sites (Fig. 5b). Additionally, Ne-CREs are relatively hypomethylated in normal prostate tissue and PRAD despite absence of H3K27 acetylation, a feature of decommissioned enhancers that were active in development (Fig. 5c)^23,24^.

**Figure 5.**
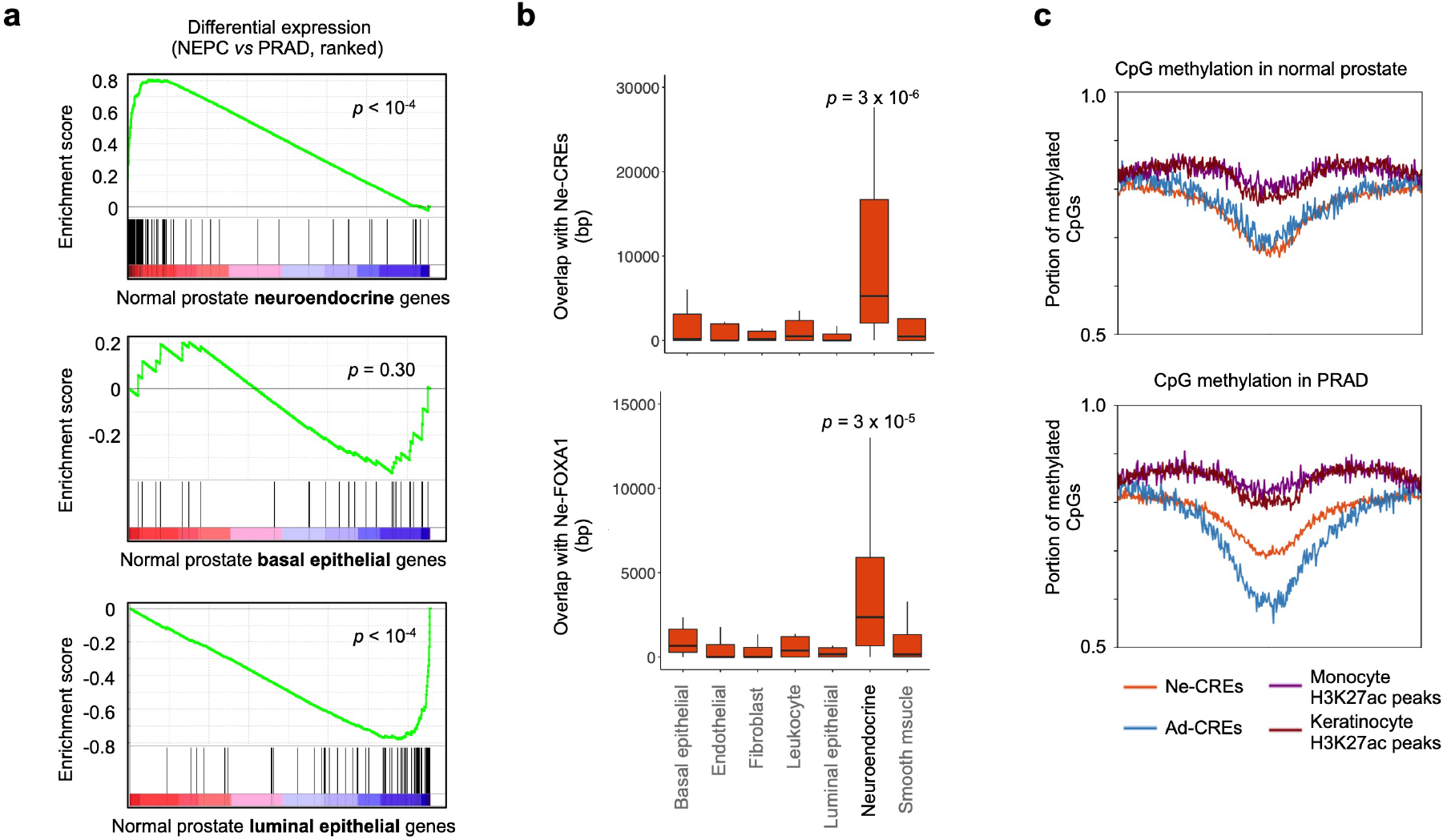
Gene expression of benign prostate cells compared to NEPC transcriptomes and epigenomes. **a,** Gene set enrichment analysis of genes specifically expressed in neuroendocrine, basal, and luminal cells from normal prostate^70^. Genes are ranked by differential expression in NEPC and PRAD LuCaP PDXs. **b**, Overlap of NEPC-enriched H3K27ac peaks (Ne-CREs; top) and FOXA1 binding sites (Ne-FOXA1; bottom) with a 200kb window centered on the transcriptional start site of the 20 most significantly differentially expressed genes in each indicated prostate cell type^70^. *p*-values correspond to Wilcoxon test of Ne-CRE/Ne-FOXA1 peak overlap near neuroendocrine cell genes versus all other indicated gene categories. **c**, fraction of CpG methylation detected by whole genome bisulfite sequencing in normal prostates tissue and PRAD at Ne-CREs and Ad-CREs. Methylation levels at H3K27ac peaks identified in epithelial keratinocytes or in peripheral blood monocytes are included for comparison. x-axis corresponds to peak center ±3kb.

We hypothesized that a neuroendocrine epigenomic program is encoded in the developmental history of the prostate, thereby priming NEPC genes for inappropriate activation under the selective pressure of androgen blockade. Consistent with this hypothesis, many genes that become highly expressed in NEPC have “bivalent” (H3K4me3^+^/H3K27me3^+^) promoter histone marks in normal prostate tissue and PRAD (Fig. 6a). Bivalent genes are thought to be poised for lineage-specific activation upon removal of H3K27me3 at the appropriate stage of development^25–27^. Our data suggested that a similar principle underlies transcriptional changes in prostate cancer lineage plasticity. H3K27me3 levels decreased in NEPC compared to PRAD at 633 gene promoters, which were enriched for binding sites of the REST repressor of neuronal lineage transcription^28^ (Supplementary Fig. 4). Similar numbers of these promoters were bivalent (H3K4me3^+^/H3K27me3^+^; n=195) and repressed (H3K4me3^-^ /H3K27me3^+^; n=229) in PRAD (Fig. 6b). Critically, however, genes with bivalent (H3K4me3^+^) promoters in PRAD became more highly expressed in NEPC (Fig. 6c) than H3K4me3^-^ genes. These bivalent genes, which included NEPC TFs *ASCL1, INSM1,* and *SOX2,* may have been prepared for activation in the development of a prostate progenitor cell. Their residual H3K4me3 and promoter hypomethylation (Fig. 6d) suggest heightened potential for re-activation^24^ in NEPC with the disruption of pro-luminal AR-driven transcriptional programs.

**Figure 6.**
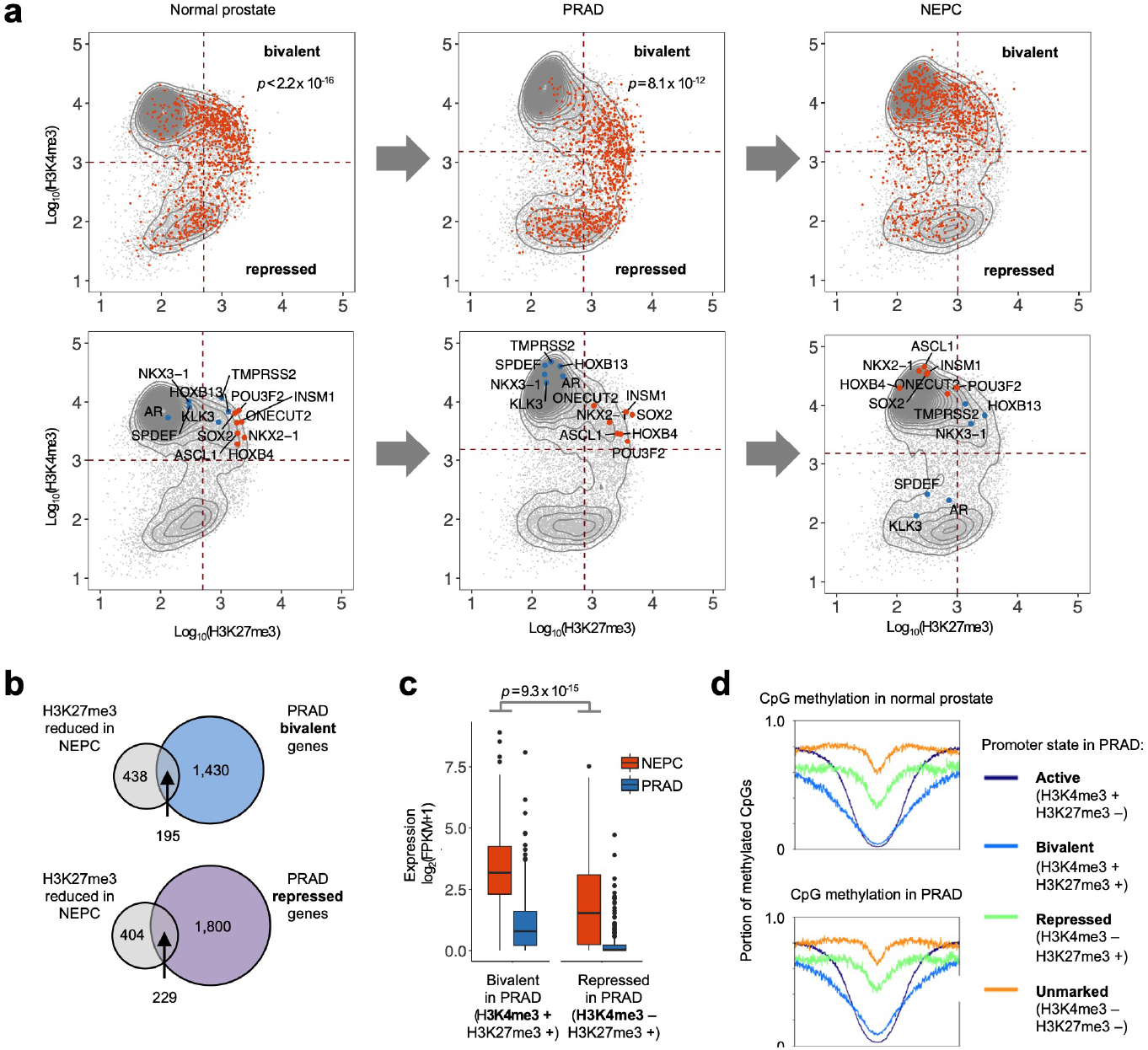
Encoding of neuroendocrine regulatory programs in the developmental history of prostate cancer. **a,** Average ChIP-seq tag density in normal prostate (n=3), PRAD (n=5) and NEPC (n=5) for H3K4me3 and H3K27me3 within 2kb of a gene transcriptional start site (TSS). Each dot represents a unique gene TSS. The top row highlights genes with upregulated expression in NEPC compared to PRAD (orange). *p*-values indicate Pearson’s Chi-squared test comparing enrichment of upregulated genes within the “bivalent” quadrant compared to the bottom two quadrants. Selected genes are highlighted in the bottom row. **b**, Intersection of genes with bivalent (H3K27me3^+^/H3K4me3^+^) or repressed (H3K27me3^+^/H3K4me3^-^) promoter annotations in PRAD and genes with reduced promoter H3K27me3 in NEPC *vs*. PRAD (log_2_ fold-change < −1, FDR-adjusted *p*-value = 0.01). **c**, Transcript expression levels in NEPC of genes whose promoters lose H3K27me3 in NEPC compared to PRAD. Genes are grouped by bivalent or repressed promoter annotations in PRAD. *p*-value corresponds to Wilcoxon ranksum test. **d**, Fraction of CpG methylation in normal prostate tissue and PRAD at TSS ± 3kb for genes in each indicated category.

## Discussion

In summary, our work demonstrates that the *cis*-regulatory landscape of prostate cancer is extensively reprogrammed in NEPC. Epigenomic profiling of human tumors identified a critical role of FOXA1 in this process, which perhaps has been overlooked because candidate drivers of NEPC have been nominated and prioritized mainly based on differential expression or somatic DNA alterations^9,11,29,30^. FOXA1 has been reported to *inhibit* neuroendocrine differentiation of prostate adenocarcinoma, based on the observations that FOXA1 is downregulated in NEPC and that FOXA1 knock-down induces neuroendocrine features in PRAD cell lines^31^. Our data demonstrate that FOXA1 remains crucial in NEPC despite consistent, modest transcript downregulation in NEPC compared to PRAD. Our H3K27ac HiChIP data reveal that in NEPC, *FOXA1* contacts distal super-enhancers that are distinct from its PRAD enhancers and contain binding sites for NE-associated TFs such as ASCL1 and INSM1 (Fig 2d. and Supplementary Fig. 5). Thus, an NEPC-specific regulatory program may maintain FOXA1 expression at lower levels that are conducive to NE gene expression, reconciling our findings with the reported pro-neuroendocrine effects of partial FOXA1 suppression in PRAD^31^. While our data show that FOXA1 is essential in NEPC, further studies are required to determine if FOXA1 cistrome reprogramming directly activates Ne-CREs and to assess its role dynamic lineage plasticity.

FOXA1 may have a more general role in controlling neuroendocrine differentiation. For example, in small cell lung cancer (SCLC), a neuroendocrine lung cancer variant that can emerge *de novo* or from *EGFR*-mutant lung adenocarcinoma after targeted kinase inhibition, *FOXA1* is highly expressed and encompassed by a super-enhancer^32^. We observe extensive H3K27 acetylation in SCLC cell lines specifically at Ne-CREs and NEPC-enriched FOXA1 binding sites, suggesting similar enhancer usage between in SCLC and NEPC (Supplementary Fig. 6), consistent with recent reports^29,33^. Ultimately, therapeutic targeting of FOXA1 and/or proteins that collaborate with or covalently modify this TF presents an attractive strategy to inhibit lineage plasticity, as FOXA1 is a common vulnerability in both PRAD and NEPC.

## Methods

### Patient-derived xenograft and tissue specimens

LuCaP patient-derived xenografts (PDXs) have been described previously^10,34–37^ with the exception of LuCaP 208.1. LuCaP 208.1 was derived from treatment-emergent NEPC and demonstrates typical small cell histology. All LuCaP PDXs were derived from resected 10 metastatic prostate cancer with informed consent of patient donors as described previously^10^ under a protocol approved by the University of Washington Human Subjects Division IRB. Liver metastasis needle biopsy specimens were obtained from the Dana-Farber Cancer Institute Gelb Center biobank and were collected under DFCI/Harvard Cancer Center IRB-approved protocols. Metastases were reviewed by a clinical pathologist. The NEPC metastasis was obtained from a patient with *de novo* metastatic prostate adenocarcinoma after 17 months of androgen deprivation therapy with leuprolide and bicalutamide. Immunohistochemistry revealed staining for synaptophysin, chromogranin, and NKX3-1 (weak), and absence of RB1, AR, and PSA.

### Epigenomic profiling

#### Histone mark ChIP in LuCaP PDXs

Chromatin immunoprecipitation (ChIP) for histone marks (H3K27ac, H3K27me3, and H3K4me3) in PDXs was performed as previously described^38^. Briefly, 20-30 mg of frozen tissue was pulverized using the CryoPREP dry impactor system (Covaris). The tissue was then fixed using 1% formaldehyde (Thermo fisher) in PBS for 18 minutes at 37 degrees Celsius and was quenched with 125 mM glycine. Chromatin was lysed in ice-cold lysis buffer (50mM Tris, 10mM EDTA, 1% SDS with protease inhibitor) and was sheared to 300~800 bp using the Covaris E220 sonicator (105 watt peak incident power, 5% duty cycle, 200 cycles/burst) for 10 min. Five volumes of dilution buffer (1% Triton X-100, 2 mM EDTA, 150 mM NaCl, 20 mM Tris HCl pH 8.1) were added to chromatin. The sample was then incubated with antibodies (H3K27ac, Diagenode, C15410196; H3K27me3, Cell Signaling 9733S; H3K4me3, Diagenode C15410003 premium) coupled with protein A and protein G beads (Life Technologies) at 4 degrees Celsius overnight. The chromatin was washed with RIPA wash buffer (100 mM Tris pH 7.5, 500 mM LiCl, 1% NP-40, 1% sodium deoxycholate) for 10 minutes six times and rinsed with TE buffer (pH 8.0) once.

#### Transcription factor ChIP in PDXs

ChIP for transcription factors (FOXA1 and ASCL1) in PDXs was performed as previously described^38^. Briefly, 50-80 mg of frozen tissue was pulverized using the CryoPREP dry impactor system (Covaris). The tissue was then fixed using 1% formaldehyde (Thermo fisher) in PBS for 18 minutes at room temperature and was quenched with 125 mM glycine. Chromatin was lysed in 1mL ice-cold Myer’s Lysis buffer (0.1% SDS, 0.5% sodium deoxycholate and 1% NP-40 with protease inhibitor) and was sheared to 300~800 bp using the Covaris E220 sonicator (140 PIP, 5% duty cycle, 200 cycles/burst) for 20 min. The sample was then incubated with antibodies (FOXA1, ab23738, Abcam; ASCL1, ab74065) coupled with protein A and protein G beads (Life Technologies) at 4 degrees Celsius overnight. The chromatin was washed with RIPA wash buffer (100 mM Tris pH 7.5, 500 mM LiCl, 1% NP-40, 1% sodium deoxycholate) for 10 minutes six times and rinsed with TE buffer (pH 8.0) once.

#### LNCaP ChIP

ChIP in LNCaP was performed as previously described^38^. 10 million cells were fixed with 1% formaldehyde at room temperature for 10 minutes and quenched. Cells were collected in lysis buffer (1% NP-40, 0.5% sodium deoxycholate, 0.1% SDS and protease inhibitor (#11873580001, Roche) in PBS)^39^. Chromatin was sonicated to 300-800 bp using a Covaris E220 sonicator (140 watt peak incident power, 5% duty cycle, 200 cycleburtst). Antibodies (FOXA1, ab23738, Abcam; H3K27ac, C15410196, Diagenode; ASCL1, ab74065) were incubated with 40 μl of Dynabeads protein A/G (Invitrogen) for at least 6 hours before immunoprecipitation of the sonicated chromatin overnight. Chromatin was washed with LiCl wash buffer (100 mM Tris pH 7.5, 500 mM LiCl, 1% NP-40, 1% sodium deoxycholate) 6 times for 10 minutes sequentially.

#### ChIP sequencing

Sequencing libraries were generated from purified IP sample DNA using the ThruPLEX-FD Prep Kit (Rubicon Genomics). Libraries were sequenced using 150-base paired end reads on an Illumina platform (Novogene).

#### ATAC-seq

LuCaP PDX tissues were resuspended and dounced in 300 ul of RSB buffer (10 mM Tris-HCl pH 7.4, 10 mM NaCl, and 3 mM MgCl2 in water) containing 0.1% NP40, 0.1% Tween-20, and 0.01% digitonin. Homogenates were transferred to a 1.5 ml microfuge tube and incubated on ice for 10 minutes. Nuclei were filtered through a 40 μm cell strainer and nuclei were washed with RSB buffer and counted. 50,000 nuclei were resuspended in 50 μl of transposition mix^40^ (2.5 μl transposase (100 nM final), 16.5 μl PBS, 0.5 μl 1% digitonin, 0.5 μl 10% Tween-20, and 5 μl water) by pipetting up and down six times. Transposition reactions were incubated at 37 C for 30 minutes in a thermomixer with shaking at 1,000 r.p.m. Reactions were cleaned with Qiagen columns. Libraries were amplified as described previously^41^ and sequenced on an Illumina Nextseq 500 with 35 base paired-end reads.

### ChIP-seq data analysis

ChIP-sequencing reads were aligned to the human genome build hg19 using the Burrows-Wheeler Aligner (BWA) version 0.7.15^42^. Non-uniquely mapping and redundant reads were discarded. MACS v2.1.1.20140616^43^ was used for ChIP-seq peak calling with a q-value (FDR) threshold of 0.01. ChIP-seq data quality was evaluated by a variety of measures, including total peak number, FrIP (fraction of reads in peak) score, number of high-confidence peaks (enriched > ten-fold over background), and percent of peak overlap with DHS peaks derived form the ENCODE project. ChIP-seq peaks were assessed for overlap with gene features and CpG islands using annotatr^44^. IGV^45^ was used to visualize normalized ChIP-seq read counts at specific genomic loci. ChIP-seq heatmaps were generated with deepTools^46^ and show normalized read counts at the peak center ± 2kb unless otherwise noted. Overlap of ChIP-seq peaks was assessed using BEDTools. Peaks were considered overlapping if they shared one or more base-pairs.

#### Identification and annotation of PRAD- and NEPC-enriched ChIP-seq peaks

Sample-sample clustering, principal component analysis, and identification of lineage-enriched peaks were performed using Mapmaker (https://bitbucket.org/cfce/mapmaker), a ChIP-seq analysis pipeline implemented with Snakemake^47^. ChIP-seq data from PRAD and NEPC LuCaP PDXs were compared to identify H3K27ac, H3K27me3, and FOXA1 peaks with significant enrichment in the NEPC or PRAD lineage. Only LuCaP PDXs from distinct patients were included, with the exception of the H3K27me3 differential peak analysis, which included both LuCaP 145.1 and 145.2, two LuCaP PDXs derived from distinct NEPC metastases from a single patient. A union set of peaks for each histone modification or TF was created using BEDTools. narrowPeak calls from MACS were used for H3K27ac and FOXA1, while broadPeak calls were used for H3K27me3. The number of unique aligned reads overlapping each peak in each sample was calculated from BAM files using BEDtools. Read counts for each peak were normalized to the total number of mapped reads for each sample. Quantile normalization was applied to this matrix of normalized read counts. Using DEseq2^48^, lineage-enriched peaks were identified at the indicated FDR-adjusted p-value (padj) and log_2_ fold-change cutoffs (H3K27ac, padj < 0.001, |log_2_ fold-change| > 3; FOXA1, padj < 0.001, |log_2_ fold-change| > 2; H3K27me3, padj < 0.01, |log_2_ fold-change| > 1). Unsupervised hierarchical clustering was performed based on Spearman correlation between samples. Principal component analysis was performed using the prcomp R function. Enriched *de novo* motifs in differential peaks were detected using HOMER version 4.7. The top non-redundant motifs were ranked by adjusted p-value.

The GREAT tool^49^ was used to asses for enrichment of Gene Ontology (GO) and MSigDB perturbation annotations among genes near differential ChIP-seq peaks, assigning each peak to the nearest gene within 500kb. The cistromedb toolkit (http://dbtoolkit.cistrome.org/) was used to compare ChIP-seq peaks for overlap with peaks from a large database of uniformly analyzed published ChIP-seq data (quantified as a “GIGGLE score”)^50^. Published TFs and histone marks were ranked by similarity to the querry dataset based on the top 1,000 peaks in each published dataset. Prior to cistromedb toolkit analysis, ChIPseq peaks were mapped from hg19 to hg38 using the UCSC liftover tool (https://genome.ucsc.edu/cgi-bin/hgLiftOver).

For analysis of H3K27 acetylation in lung cancer at lineage-enriched candidate regulatory elements, fastq files were generated from sequence read archives (SRA) from published ChIP-seq experiments for SCLC^51^ and LUAD^52–55^ (SRA numbers SRR568435, SRR3098556, SRR4449027, SRR4449025, and SRR6124068).

For Fig. 5c, H3K27ac ChIP-seq peaks from primary peripheral blood monocytes (ENCFF540CVX) and epithelial keratinocytes (ENCFF943CBQ)^53^ were used as a comparator to peaks derived from LuCaP PDXs. For these comparisons, monocyte and keratinocyte peaks within 1kb of a LuCaP peak were excluded.

### RNA-seq and differential expression analysis

RNA-seq data from human adenocarcinoma and NEPC have been reported previously^8^ and were obtained from dbGaP (accession number phs000909.v1.p1). Transcriptomes were sequenced from two replicates from each of five PRAD LuCaP PDXs (23, 77, 78, 81, and 96) and five NEPC LuCaP PDXs (49, 93, 145.1, 145.2, and 173.1). RNA concentration, purity, and integrity were assessed by NanoDrop (Thermo Fisher Scientific Inc.) and Agilent Bioanalyzer. RNA-seq libraries were constructed from 1 μg total RNA using the Illumina TruSeq Stranded mRNA LT Sample Prep Kit according to the manufacturer’s protocol. Barcoded libraries were pooled and sequenced on the Illumina HiSeq 2500 generating 50 bp paired end reads. FASTQ files were processed using the VIPER workflow^56^. Read alignment to human genome build hg19 was performed with STAR^57^. Cufflinks was used to assemble transcript-level expression data from filtered alignments^58^. Differential gene expression analysis (NEPC *vs*. PRAD) was conducted using DESeq2^48^.

### H3K27ac HiChIP

Pulverized frozen tissue from LuCaP 173.1 was fixed with 1% formaldehyde in PBS at room temperature for 10 minutes as previously described^38^. Sample was incubated in lysis buffer and digested with MboI (NEB) for 4 hours. After 1 hour of biotin incorporation with biotin dATP, the sample was ligated using T4 DNA ligase for 4 hours. Chromatin was sheared using 140 PIP, 5% duty cycle, and 200 cycles/burst for 8 minutes in shearing buffer composed of 1% NP-40, 0.5% sodium deoxycholate, and 0.1% SDS in PBS (LNCaP) or using 100 PIP, 5% duty cycle, 200 cycles/burst for 3 minutes in 1% SDS, 50mM Tris (pH 8.1), and 5mM EDTA (LuCaP 173.1). ChIP was then performed using H3K27Ac antibody (Diagenode, C1541019)^59^.

Immunoprecipitated sample was pulled down with streptavidin C1 beads (Life Technologies) and treated with Transposase (Illumina). Amplification was performed for the number of cycles required to reach 1/3 of the maximal fluorescence on qPCR plot with SYBR^®^ Green I(Life Technologies). Libraries were sequenced using 150-base paired end reads on the Illumina platform (Novogene).

#### Alignment and filtering using HiC-Pro

We processed paired-end fastq files using HiC-Pro^60^ to generate intra-and inter-chromosomal contact maps. The reads were first trimmed to remove adaptor sequences using Trim Galore *(https://github.com/FelixKrueger/TrimGalore).* Default settings from HiC-Pro were used to align reads to the hg19 human genome, assign reads to MboI restriction fragments, and remove duplicate reads. Only uniquely mapped valid read pairs involving two different restriction fragments were used to build the contact maps.

#### FitHiChIP

We applied FitHiChIP^61^ for bias-corrected peak calling and DNA loop calling. We used MACS2 broadPeak peak calls from H3K27ac ChIP-seq in LuCaP 173.1 (NEPC). 44,609 peaks were called at a q-value < 0.01. We used a 5Kb resolution and considered only interactions between 5kb-3Mb. We used peak-to-peak (stringent) interactions for the global background estimation of expected counts (and contact probabilities for each genomic distance), and peak-to-all interactions for the foreground, meaning at least one anchor must overlap a H3K27ac peak. The corresponding FitHiChiP options specified are “IntType=3” and “UseP2PBackgrnd=1”.

#### Assignment of enhancer-promoter interactions using H3K27ac HiChIP data

NCBI RefSeq genes (hg19) were downloaded from the UCSC genome table browser (https://genome.ucsc.edu/cgi-bin/hgTables). Only uniquely mapping genes were considered. The longest transcript was selected for genes with multiple annotated transcripts. We searched for H3K27ac HiChIP loops with one anchor (defined with a 5kb window) overlapping a region between 0 and 5kb upstream of a gene transcriptional start site. We selected subset of these loops for which the second anchor (with a 5kb window) overlapped with H3K27ac peaks identified by ChIP-seq in LuCaP 173.1 (NEPC) or with NEPC-enriched H3K27ac peaks (Ne-CREs). Gene promoters and distal H3K27ac peaks / Ne-CREs were considered looped if each overlapped with an anchor of the same high-confidence H3K27ac HiChIP loop(s). To examine the association of regulatory element looping with gene expression, genes were binned by the number of distinct, looped Ne-CREs or H3K27ac peaks. Differential expression between NEPC and PRAD LuCaP PDXs, as assessed by DESeq2 analysis of LuCaP RNA-seq data, was plotted for genes in each bin. Wilcoxon rank-sum *p*-values were calculated for differential expression of genes looped to one versus two or more H3K27ac/Ne-CRE peaks. A *p*-value < 0.01 was considered significant.

### Master transcription factor analysis

#### Super-enhancer ranking analyses

Enhancer and super-enhancer (SE) calls were obtained using the Rank Ordering of Super-enhancer (ROSE2) algorithm^12^. We selected SEs assigned to transcription factors (TFs)^62,63^, and for each sample, we obtained the ranks of all TF SEs. Considering only the top 5% TFs by median ranking in NEPC or PRAD, we applied a one-sided Mann–Whitney U test to identify lineage-enriched TF SEs (FDR = 10%).

#### Clique enrichment and clustering analysis

Clique enrichment scores (CESs) for each TF were calculated using clique assignments from Coltron^64^. Coltron assembles transcriptional regulatory networks (cliques) based on H3K27 acetylation and TF binding motif analysis. The clique enrichment score for a given TF is the number of cliques containing the TF divided by the total number of cliques. We incorporated ATAC-seq data to restrict the motif search to regions of open chromatin. Using the CES, we performed clustering (distance = Canberra, agglomeration method = ward.D2) considering only TFs that appear in cliques in at least 80% of the samples in at least one lineage group (4 out of 5 NEPC and 11 out of 14 PRAD).

#### Motif enrichment at super-enhancers with loops to the *FOXA1* locus

H3K27ac HiChIP data were used to select distal SEs that form three-dimensional contacts with the *FOXA1* locus. We used the Coltron algorithm to search for TF motifs in ATAC-seq peaks within these SEs. We considered all TFs that were categorized as expressed by Coltron based on H3K27ac levels at the TF gene locus. Motif enrichment for a TF was calculated as the total number of non-overlapping base pairs (bp) covered by the TF motif, divided by the summed length (in bp) of the SEs. Values in the heatmap legend correspond to percent coverage (i.e., the largest value corresponds to 0.4%).

### FOXA1 mutational profiling

*FOXA1* mutational status was assessed from exome sequence data (62x-110X depth of coverage). Each LuCaP PDX was sequenced using the Illumina Hi-seq platform with 100 bp paired-end reads. Hybrid capture was performed SeqCapV3. Mouse genome subtraction was performed using the mm10 genome build and reads were aligned to human reference genome hg19. For sequence analysis, bam files processed as per Genome Analysis Toolkit (GATK) best practice guideline^65^. Mutation pathogenicity was annotated using Clinvar, OncoKb and Civic. We Used MuTect1 and Unified Genotyper for mutation calls. Copy number was derived using the Sequenza R package.

### FOXA1 siRNA knock-down

WCM154 organoids were cultured and maintained as previously described^19^. Organoids were dissociated to single cells using TrypLE (ThermoFisher). One million cells were resuspended in 20μl of electroporation buffer (BTXpress) and mixed with 60 pmole of control or FOXA1 On-target pool siRNA (Dharmacon). Then organoid-siRNA mixtures were transferred to a 16-well NucleocuvetteTM Strip and nucleofection was performed in a 4D-Nucleofector (Lonza). Following nucleofection, 10^5^ organoids cells were grown in a 12-well plate coated with 1% collagen I (ThermoFisher) for 7 days. Both adherent and floating cells were collected and stained with 0.4% trypan blue solution (ThermoFisher). Total cell numbers were measured by a hemocytometer. Cell proliferation with FOXA1 knock-down was normalized to control siRNA cells.

### FOXA1 shRNA knock-down

LNCaP, LNCaP 42D, and LNCaP 42F cells were seeded in parallel 6-well plates at 500k, 500k, or 100k, respectively. 24 hours later, cells were infected with lentivirus containing shRNAs targeting GFP control or *FOXA1.* 48 hours following infection, equal cell numbers were seeded, and proliferation was assayed 6 days later using a Vi-Cell. 72 hours following infection, a second plate infected in parallel was harvested for immunoblotting. The target sequence against GFP was CCACATGAAGCAGCACGACTT (shGFP). The target sequences against FOXA1 were GCGTACTACCAAGGTGTGTAT (shFOXA1-1) and TCTAGTTTGTGGAGGGTTAT (shFOXA1-2).

### FOXA1 CRISPR-Cas9 knock-out

Blasticidin-resistant Cas9 positive LNCaP, LNCaP 42D, and LNCaP 42F cells were cultured in 20μg/mL blasticidin (Thermo Fisher Scientific, NC9016621) for 72 hours to select for cells with optimal Cas9 activity. LNCaP, LNCaP 42D, and LNCaP 42F, PC3M cells were seeded in parallel 6-well plates at 300k, 300k, 300k, or 60k, respectively. Cells were infected after 24 hours with lentiviruses expressing sgRNAs targeting GFP control or *FOXA1.* Cells were subject to puromycin selection and harvested for immunoblot after 3 days. 6 days following selection, cell viability was determined using a Vi-Cell. The target sequences against GFP were AGCTGGACGGCGACGTAAA (sgGFP1) and GCCACAAGTTCAGCGTGTCG (sgGFP2). The target sequences against FOXA1 were GTTGGACGGCGCGTACGCCA (sgFOXA1-1), GTAGTAGCTGTTCCAGTCGC (sgFOXA1-2), CAGCTACTACGCAGACACGC (sgFOXA1-3), and ACTGCGCCCCCCATAAGCTC (sgFOXA1-4).

### Western Blots

For WCM154 Western blots, cell pellets were lysed in RIPA buffer (MilliporeSigma, 20-188) supplemented with Protease/Phosphatase Inhibitor Cocktail (Cell Signaling Technology, 5872S). Protein concentrations were assayed with a Pierce BCA Protein Assay Kit (Thermo Fisher Scientific, PI23225), and protein was subsequently denatured in NuPAGE LDS sample buffer (Thermo Fisher Scientific, NP0007) containing 5% β-Mercaptoethanol. 13μg of each protein sample was loaded onto NuPAGE 4-12% Bis-Tris Protein gels (Thermo Fisher Scientific), and samples were run in NuPAGE MOPS SDS Running Buffer (Thermo Fisher Scientific, NP0001). Following electrophoresis, proteins were transferred to nitrocellulose membranes via an iBlot apparatus (Thermo Fisher Scientific). After blocking in Odyssey Blocking Buffer (LI-COR Biosciences, 927-70010) for one hour at room temperature, membranes were cut and incubated in primary antibodies diluted 1:1000 in Odyssey Blocking Buffer overnight at 4°C. The next morning, membranes were washed three times with Phosphate-Buffer Saline, 0.1% Tween (PBST) and then incubated with fluorescent anti-rabbit secondary antibodies (Thermo Fisher Scientific, NC9401842) for one hour at room temperature. Membranes underwent five PBST washes and were then imaged using an Odyssey Imaging System (LI-COR Biosciences). Primary antibodies used include FOXA1 (Cell Signaling Technology, 58613S) and β-actin (Cell Signaling Technology, 8457L).

For LNCaP, LNCaP 42D, and LNCaP 42F Western Blots, cell lysate was extracted using RIPA lysis buffer (Sigma) containing protease inhibitor (Roche) and phosphatase inhibitor (ThermoFisher). 50 μg of protein was subjected to a 4-15% Mini-PROTEAN Precast electrophoresis gel (Bio-Rad) then transferred to 0.22 um nitrocellulose membrane (Bio-Rad) and blocked in 5% blotting grade blocker (Bio-Rad). Membranes were incubated with primary antibodies overnight (FOXA1, Abcam, 1:2000, ab23738; Synaptophysin, Cell Marque, 1:5000, MRQ-40; INSM1, Santa Cruz, 1:2000, sc-377428; FOXA2, Abcam; 1:2500, ab108422; Chromogranin A, Abcam, 1:2000, ab15160; Vinculin Cell signaling, 1:5000, #13901). Membranes were then washed in 1x Tris-buffered saline with 0.5% Tween-20 (Boston BioProducts) and incubated with secondary antibodies (mouse, Bio-Rad, 1:2500; rabbit, BioRad, 1:2500). Western HRP substrate kit was used to detect chemiluminescent signal (Millipore, Classico).

### Analysis of FOXA1 binding sites across prostate cancer states

FOXA1 cistromes were compared across different states of prostate cancer progression (normal prostate, prostate-localized adenocarcinoma, PDXs derived from metastatic castration resistant prostate cancer, and PDXs derived from NEPC). FOXA1 ChIP from normal prostate tissue and prostate-localized adenocarcinoma will be reported separately (Pomerantz *et al.,* submitted). For normal prostate tissue FOXA1 ChIP, tissue cores were obtained from regions of prostatectomy specimens with dense epithelium and no evidence of neoplasia on review by a genitourinary pathologist. PDX samples used are listed in Table S1. PDXs derived from localized prostate cancer were excluded from this analysis. Because the normal prostate and localized adenocarcinoma samples were sequenced with single-end sequencing with an average of ~20 million reads, paired-end sequencing data from LuCaP PDXs were downsampled to 20M reads, using a single end trimmed to 75 base-pairs using seqtk (https://github.com/lh3/seqtk).

Pairwise comparisons were made between normal prostate (N=5) and localized PRAD (N=5), localized PRAD and metastatic PRAD PDXs (N=11), and metastatic PRAD PDXs and NEPC PDXs (N=5) using DESeq2 as described above. Peaks were considered significantly different between groups at a log2 |fold-change| threshold of 2 and FDR-adjusted *p*-value threshold of 0.001. “Shared” peaks were defined as the intersection of all peaks that were present in each group but not significantly different in any comparison.

### Immunohistochemistry

Immunohistochemistry was performed on tissue microarray (TMA) sections. TMA slides were stained for FOXA1 (Abcam ab170933, 1:100 dilution with 10 mM NaCitrate antigen retrieval) and FOXA2 (Abcam ab108422, 1:500 dilution with 10 mM NaCitrate antigen retrieval) using a standard procedure^66^. Rabbit IgG was used as a negative control. Nuclear staining intensity was assigned levels 0, 1+, 2+, or 3+ and H-scores were calculated as: [1 x (% of 1 + cells) + 2 x (% of 2+ cells) + 3 x (% of 3+ cells)]. Evaluations were performed in a blinded fashion.

### ASCL1/NKX2-1 overexpression in LNCaP

#### Transduction of LNCaP cells with ASCL1 and NKX2-1

The open reading frames of ASCL1 and NKX2-1 were cloned into the pLX_TRC302 lentiviral expression vector (Broad Institute) using the gateway recombination system. A construct expressing eGFP (pLX_TRC302_GFP) was used as a negative control. Viruses were generated by transfecting 293T cells with packaging vectors pVsVg and pDelta8.9. Supernatant was collected after 48 hours. LNCaP cells were transduced in the presence of 4μg/ml polybrene and harvested after 3 days for RNA-seq, ATAC-seq, and ChIP-seq.

ChIP seq was performed as described above, using 10-15 million cells fixed with 1% paraformaldehyde for 10 minutes at room temperature, followed by quenching with glycine. RNA was isolated using QIAGEN RNeasy Plus Kit and cDNA synthesized using Clontech RT Advantage Kit. Quantitative PCR was performed on a Quantstudio 6 using SYBR green. The following primers were used for qRT-PCR:

AR qRT-PCR fwd GTGTCAAAAGCGAAATGGGC
AR qRT-PCR rev GCTTCATCTCCACAGATCAGG
ASCL1 qRT-PCR fwd CTACTCCAACGACTTGAACTCC
ASCL1 qRT-PCR rev AGTTGGTGAAGTCGAGAAGC
GAPDH qRT-PCR fwd CATGAGAAGTATGACAACAGCCT
GAPDH qRT-PCR rev AGTCCTTCCACGATACCAAAGT
SOX2 qRT-PCR fwd CACACTGCCCCTCTCAC
SOX2 qRT-PCR rev TCCATGCTGTTTCTTACTCTCC
SYP qRT-PCR fwd AGACAGGGAACACATGCAAG
SYP qRT-PCR rev TCTCCTTAAACACGAACCACAG

### Analysis of promoter H3K4 and H3K27 trimethylation

Refseq gene coordinates (hg19) were compiled, selecting the longest isoform where multiple were annotated. Normalized tag counts from H3K27me3 and H3K4me3 ChIP-seq within 2kb of each transcriptional start site (TSS) were calculated for each sample, then averaged across multiple samples in each group (five NEPC PDXs, five PRAD PDXs, three normal prostates; Pomerantz et al., submitted). Contours were calculated using the R function geom_density_2d from the ggplot2 package; they represent the 2d kernel density estimation for all included transcriptional start sites. Gene promoters were assigned “active”, “bivalent”, “unmarked”, and “repressed” annotations based on H3K4me3 and H3K27me3 levels. High/low cutoffs for these marks were determined as follows. First, the H3K4me3 normalized tag counts near each TSS were fit to two normal distributions using the normalmixEM R function from the mixtools R package. The cutoff between H3K4me3-high and -low was set at four standard deviations below the mean value of the H3K4me3-high distribution. Next, the normalized H3K27me3 tag counts near H3K4me3-high TSSs were fit to two normal distributions. The cutoff for H3K27me3-high promoters was set at four standard deviations above the mean value of the H3K27me3-low distribution. The Pearson Chi-squared test was used to quantify significance of enrichment of NEPC-upregulated genes in the “bivalent” quadrant compared to “repressed” or “unmarked” quadrants. NEPC-upregulated genes were defined as those with log_2_ fold-change > 3 and adjusted *p*-value < 1 x 10^-6^ in NEPC *vs*. PRAD. The results of the analysis were robust to using other *p*-value and differential expression thresholds.

### Methylation analysis of normal prostate

Whole genome bisulfite sequencing data from histologically normal prostate tissue were reported previously^67^ and processed as previously described^68^. CpG methylation at indicated sites was visualized using deepTools^46^.

## Data Availability

Sequence data in fastq format from this study will be deposited in GEO. Requests for LuCaP PDXs should be directed to Dr. Eva Corey

## Acknowledgements

This work was supported by the PNW Prostate Cancer SPORE P50 CA097186, DOD W81XWH-17-1-0415, P01 CA163227, R01 CA233863, The Prostate Cancer Foundation, The Richard M. Lucas Foundation, the European Union’s Horizon 2020 Research and Innovation programme under the Marie Skłodowska-Curie grant agreement No. 754490, the National Cancer Institute (T32CA009172), and by Rebecca and Nathan Milikowsky. We would like to thank the patients who generously donated tissue that made this research possible.

## Author contributions

S.C.B. analyzed ChIP-seq data and wrote the manuscript. X.Q. assisted with ChIP-seq data analysis. D.Y.T., J.H., and T.A. performed ASCL1/NKX2-1 overexpression experiments. S.Y.K. performed FOXA1 siRNA experiments under supervision of H.B. J.H., T.A., R.A., and S.A. performed FOXA1 shRNA and CRISPR experiments. E.O., C.B., and S.A.A. performed ChIP-seq experiments. J.-H.S. performed HiChIP experiments. C.G. and B.P analyzed HiChIP data. R.I.C. and M.A.S.F. performed core regulatory analysis under supervision of K.L. P.C. and K.L. performed ATAC-seq under supervision of H.W.L. and M.B. M.H. and A.N. assisted with procurement of clinical samples. J.E.B. and K. K. assisted with analysis of WGBS methylation data. L.B. performed immunohistochemistry experiments. I.M.C., and A.K. performed RNA-seq under the supervision of P.S.N. H.H.N., C.M., and E.C. provided LuCaP PDXs. E.C., M.M.P. and M.L.F. supervised the project.

## Competing interests statement

W.C.H. is a consultant for Thermo Fisher, Solasta Ventures, iTeos, Frontier Medicines, Tyra Biosciences, MPM Capital, KSQ Therapeutics, and Paraxel and is a founder of KSQ Therapeutics.

## Supplementary Figures

**Supplementary Figure 1.**
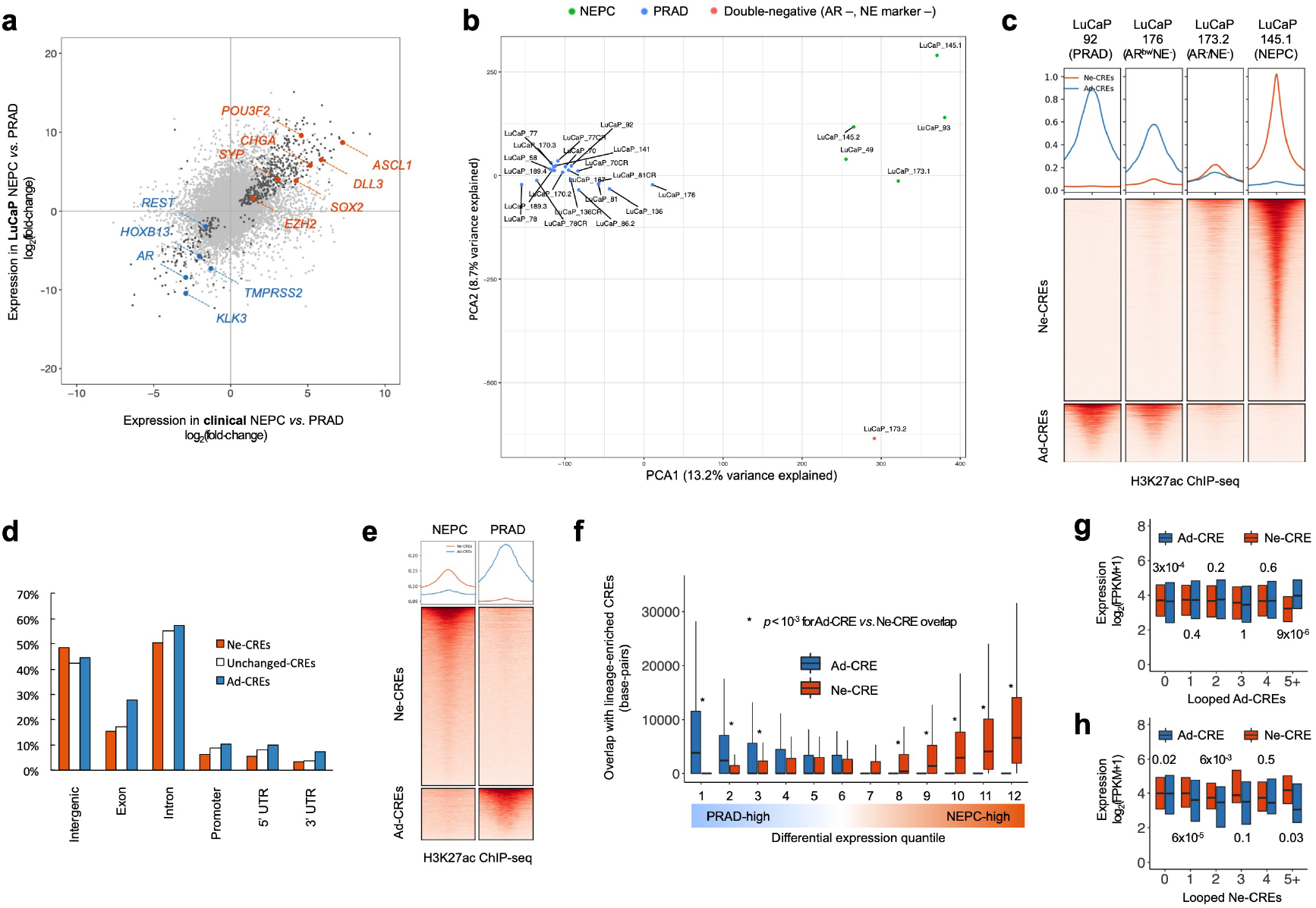
Epigenomic divergence of NEPC and PRAD. **a**, Comparison of differential gene expression between PRAD and NEPC in LuCaP PDXs (5 of each lineage, with two replicates for each sample) and clinical prostate tumors^8^. Dark gray signifies genes with significant differential expression (p<10^-4^) in both PDXs and clinical tumors. **b**, Principal component analysis of PRAD and NEPC PDXs based on H3K27ac profiles. “DN” indicates a “double-negative” PDX lacking AR or NE marker expression. **c**, Normalized H3K27ac tag density for AR^-^/NE^-^ and AR^low^/NE^-^ PDX at Ne-CREs and Ad-CREs, compared to representative PRAD and NEPC PDXs. Profile plots (top) indicate the average tag density at Ne-CREs (orange) and Ad-CREs (blue). **d,** Genomic annotations for lineage-specific and shared H3K27ac peaks. **e**, Normalized H3K27ac tag density at Ne-CREs and Ad-CREs in a clinical NEPC liver metastasis and a PRAD liver metastasis. Profile plots (top) indicate the average tag density at Ne-CREs (orange) and Ad-CREs (blue). **f**, Overlap of Ne-CREs and Ad-CREs with a 200kb window centered around the transcriptional start sites of differentially expressed genes. **g-h**, expression of genes with the indicated number of distinct looped Ad-CREs (g) or Ne-CREs (h) detected by H3K27ac HiChIP in LuCaP 173.1 (NEPC). All *p*-values were derived from Wilcoxon paired samples tests.

**Supplementary Figure 2.**
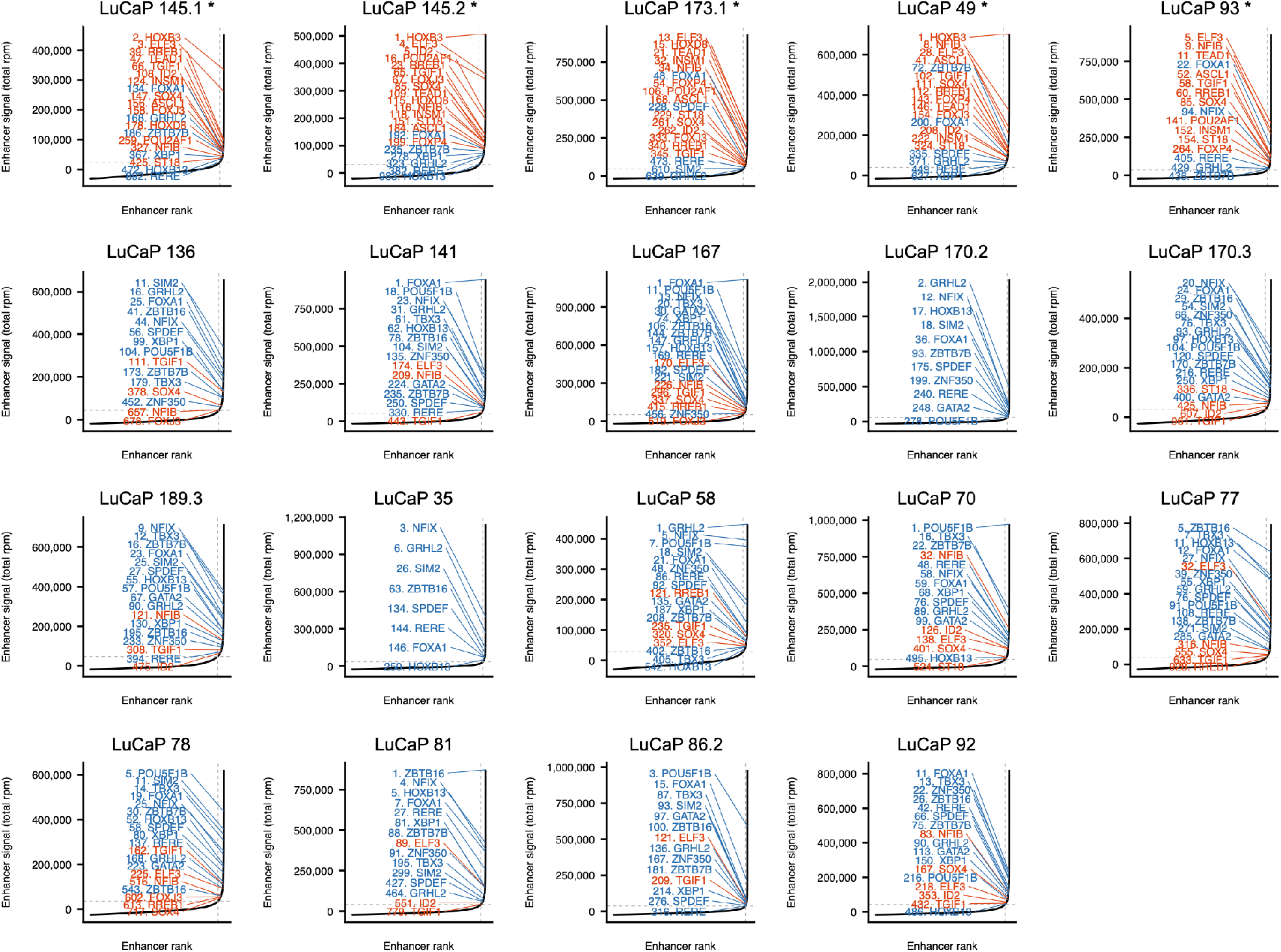
Super-enhancers (SEs) encompassing transcription factor genes that are differentially H3K27 acetylated in NEPC *vs.* PRAD. SEs are ranked by H3K27ac signal. NEPC-enriched SEs are shown in orange; PRAD-enriched SEs are shown in blue (methods). Asterisk (*) indicates NEPC LuCaP PDXs.

**Supplementary Figure 3.**
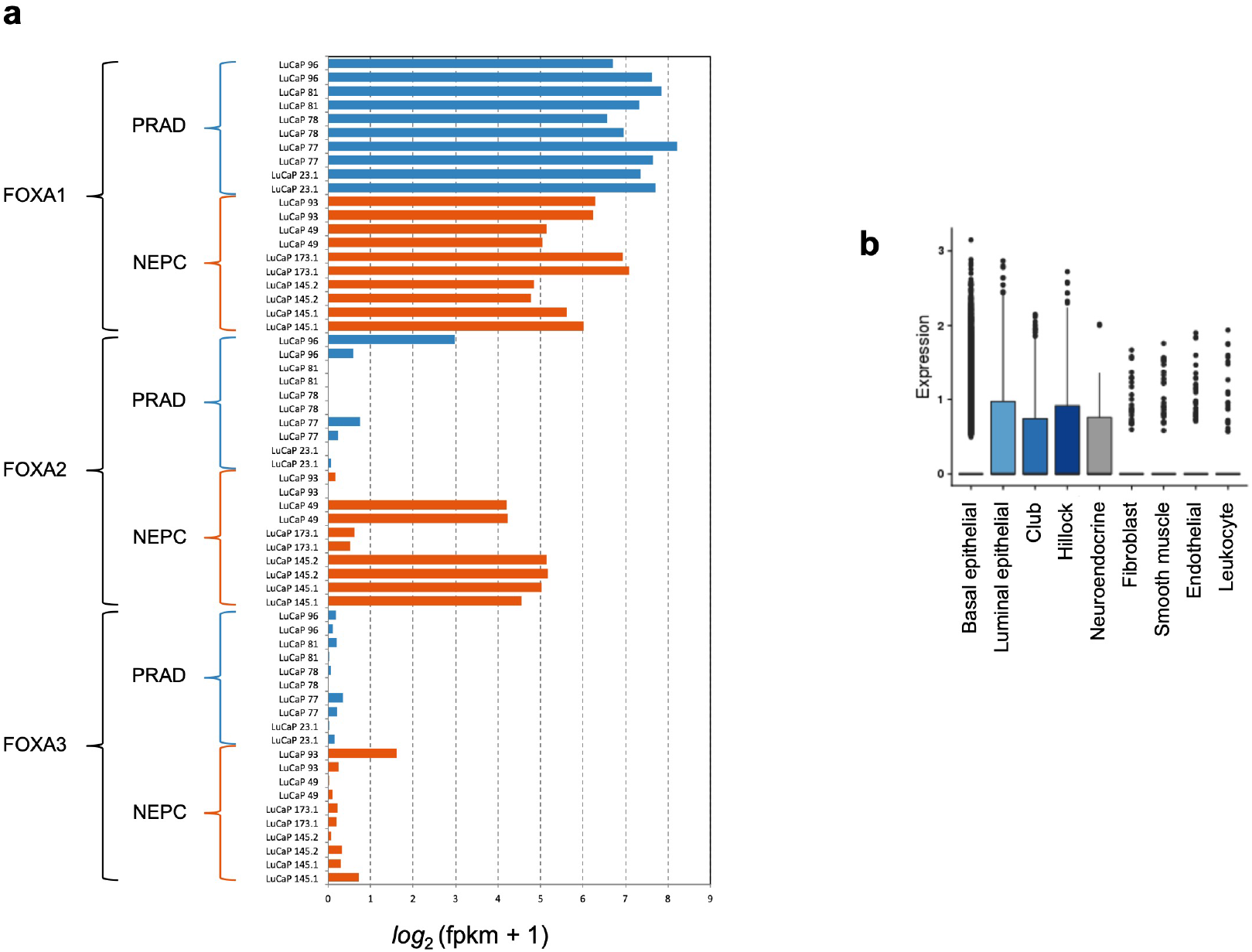
Expression of FOXA1 in PRAD, NEPC, and benign prostatic tissue. **a**, Transcript expression of FOXA family members in 5 NEPC and 5 PRAD LuCaP PDXs (two replicates each) by RNA-seq. **b**, FOXA1 expression across benign prostate cell types in a published single-cell transcriptome sequencing dataset^70^. Boxes represent the interquartile range.

**Supplementary Figure 4.**
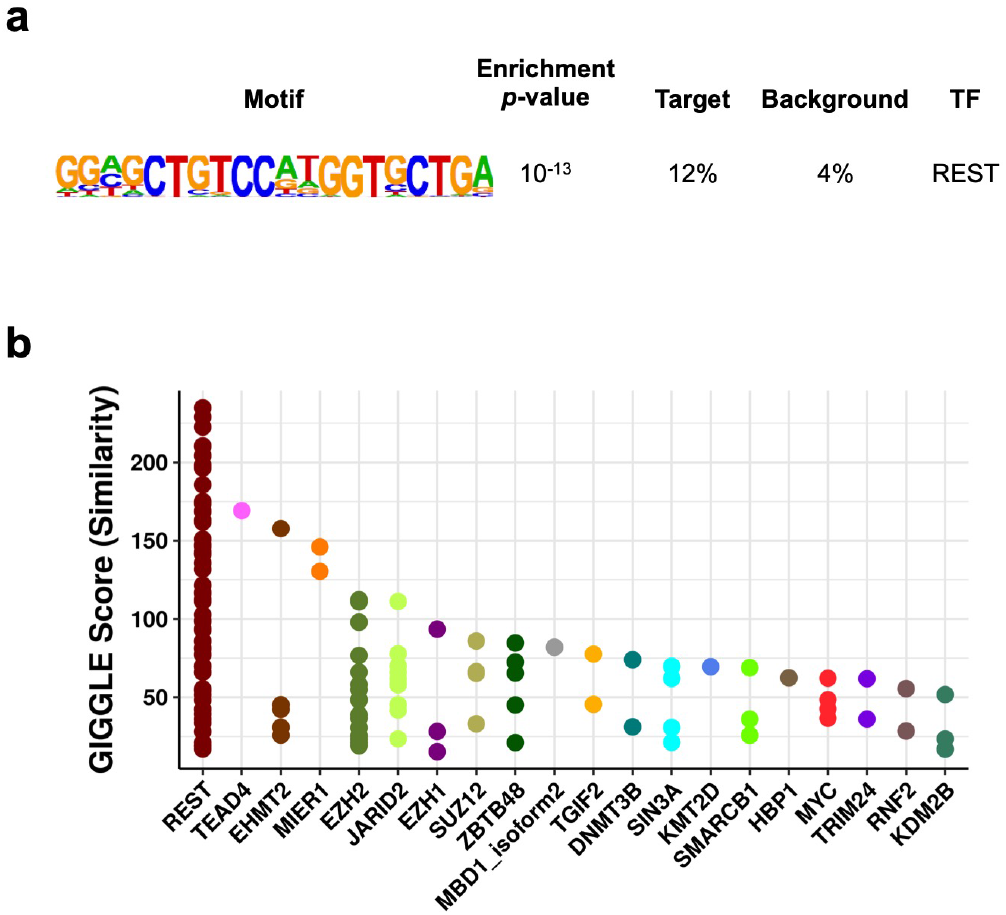
Motif enrichment and TF binding of differentially H3K27 trimethylated promoters. **a,** Motif enrichment of promoters with diminished H3K27me3 in NEPC compared to PRAD. Only the indicated motif was significantly enriched. **b**, Cistromedb toolkit analysis of published ChIP-seq datasets (dbtoolkit.cistrome.org), ranked by their degree of overlap with the differentially H3K27 trimethylated promoters analyzed in **a**.

**Supplementary Figure 5.**
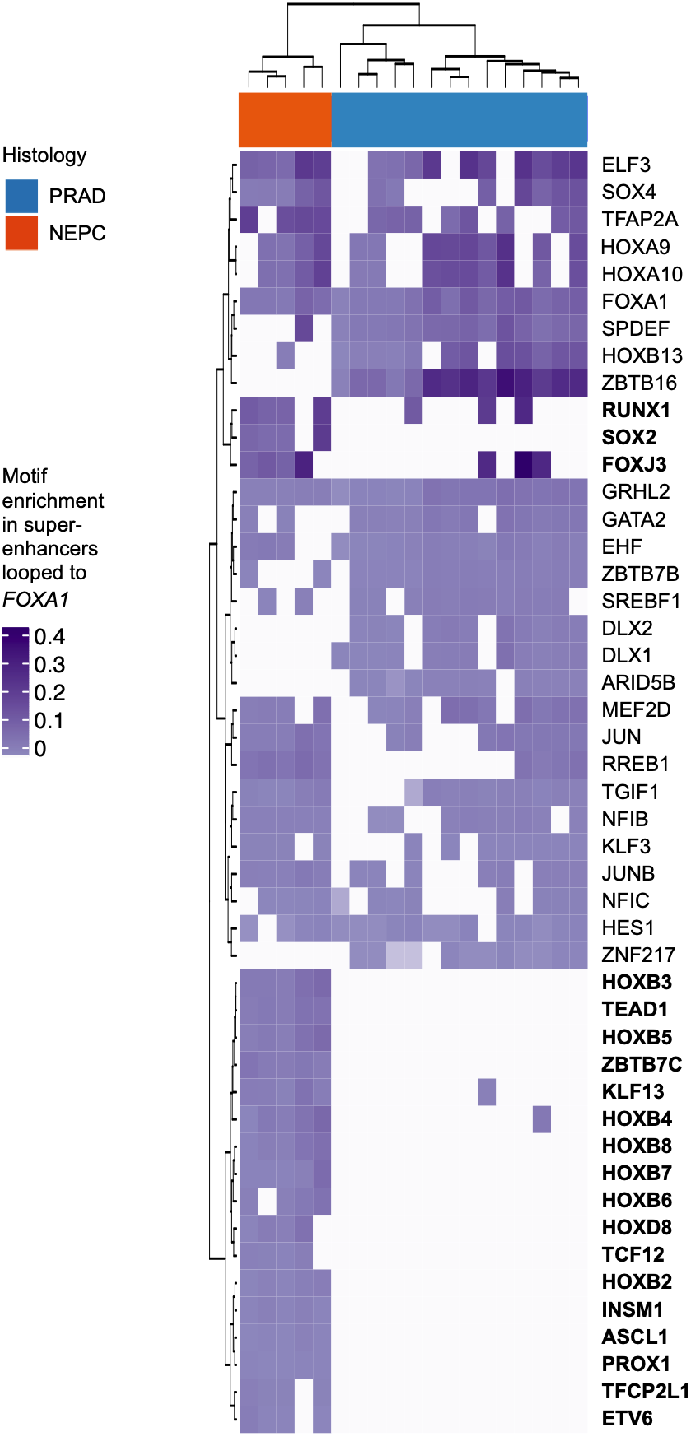
Candidate TFs involved in regulation of the FOXA1 locus. Normalized abundance of TF binding motifs at superenhancers looped to the *FOXA1* locus, as assessed by H3K27ac Hi-ChIP in NEPC (LuCaP 173.1) and PRAD (LNCaP). TFs with motif enrichment primarily in NEPC are shown in bold.

**Supplementary Figure 6.**
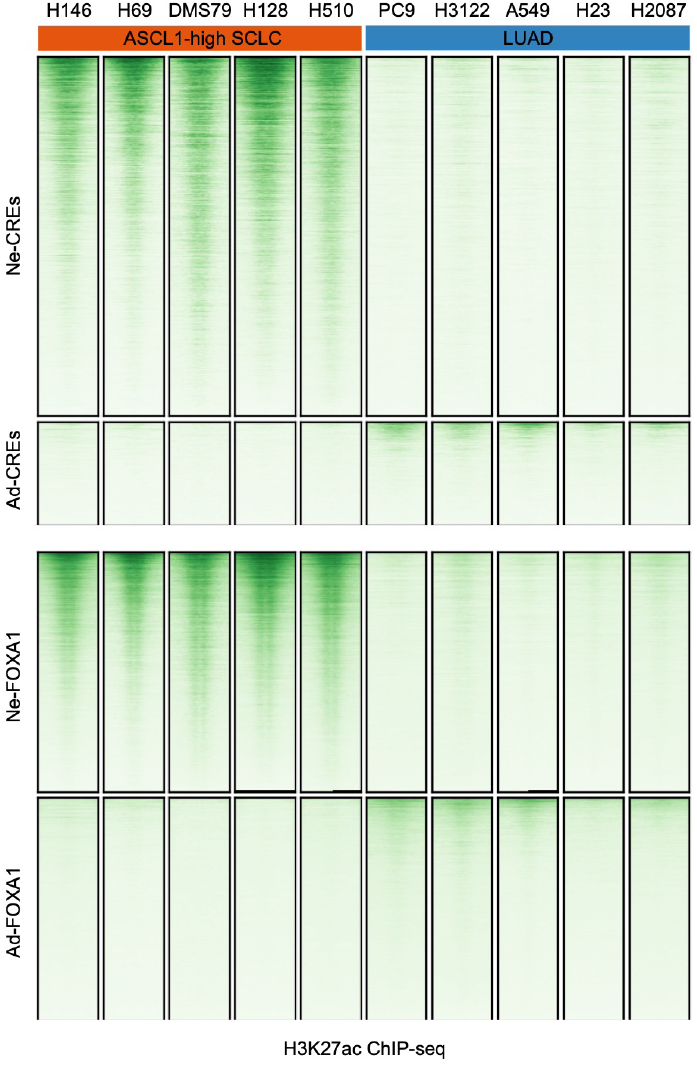
Activation of neuroendocrine candidate regulatory elements in small cell lung cancer. H3K27ac ChIP-seq profiles of ASCL1-high small cell lung cancer (SCLC) cell lines^51^ at Ne-CREs and Ad-CREs (top) and at NEPC-enriched and PRAD-enriched FOXA1 binding sites (bottom). Five lung adenocarcinoma (LUAD) cell lines are shown for comparison.

